# Parallel trophic diversifications in polyploid cyprinid fish from East Africa: from preadaptive polymorphism to trophic specialization

**DOI:** 10.1101/2023.08.18.553843

**Authors:** Boris A. Levin, Aleksandra S. Komarova, Alexei V. Tiunov, Alexander S. Golubtsov

## Abstract

Trophic diversification is one of the main mechanisms driving the adaptive radiation. The polyploid lineage of the cyprinid genus *Labeobarbus* represent an excellent model for studying the trophically-based adaptive radiation in either lacustrine or riverine environments. Recently discovered four diversifications in rivers of the Ethiopian Highlands (East Africa) demonstrate independently evolved repeated mouth polymorphisms each represented by four core mouth phenotypes: (i) generalized, (ii) thick-lipped, (iii) scraping, and iv) large-mouthed. Mouth phenotypes in some radiations can be further divided to subtypes representing from four to eight sympatric ecomorphs. Using the stable isotope and gut content analyses we tested hypothesis on trophic resource partitioning within each radiation, revealed disparity in degree of diversification between radiations and tried to reconstruct the process of trophic diversification. Three of four radiations demonstrated partitioning of trophic resources within five trophic niches: i) detritophagy, ii) macrophytophagy, iii) invertivorous benthophagy, iv) periphyton feeding, and v) piscivory. The studied riverine radiations were likely at the different stages of the diversification. One radiation having a similar set of mouth phenotypes was not trophically divergent displaying a remarkable decouple of form and function. A unique case of ecologically non-functional mouth polymorphism at an incipient stage of trophic diversification supports a concept of the plasticity-first evolution. This phenomenon stems from the pre-existing genomic templates of mouth polymorphism ancestrally inherited upon the allopolyploid origin of the *Labeobarbus* lineage. The predetermined and preadaptive mouth polymorphism can be considered a key innovation of the *Labeobarbus* that promoted to resource-based diversification via adaptive radiation.

## 1. Introduction

Trophic divergence is one of the main drivers of ecological speciation and adaptive radiation (Schluter, 2000; Seehausen & Wagner, 2014; Martin & Richards, 2019). The fishes being the most diverse group among vertebrates display numerous examples of trophic diversification during adaptive radiation - not only among textbook examples from cichlids, whitefish, Arctic charrs (Seehausen & Wagner, 2014) - but also in other taxa. Despite tremendous amount of studies focused on the mechanisms underpinning the rapid diversification during adaptive radiation (and achieved progress – e.g., Brawand *et al*., 2014; McGee *et al*., 2020; Ronco *et al*., 2021), the phenomenon is still poorly understood (Martin & Richards, 2019; Gillespie *et al*., 2020). One of the greatest findings of the last decade – a contribution of the heterogenous genome (usually evolved via past hybridization) into accelerated rates of ecological diversification and speciation (Nosil *et al*., 2009; Genner & Turner, 2012; Meier *et al*., 2017; Irisarri *et al*., 2018; Marques *et al*., 2019; Svardal *et al*., 2020). One of the mechanisms to increase genetic heterogeneity is a polyploidization. After a round of fish-specific genome duplication (Meyer & Van de Peer, 2005) occurred ca. 350 mya in the ray-finned fish (Actinopterygii) lineage, the numerous further polyploidization events happened within and resulted in >1000 species mostly within Cypriniformes, Salmoniformes and Acipenseriformes (Yang *et al*., 2022). Contrary to plants, the animal polyploidization is realized predominantly *via* allopolyploidization (Gregory & Mable, 2005) that means the most of fish polyploid lineages contains the different subgenomes. Therefore, the polyploid lineages look as perfect candidates to test a hypothesis on role of complex genome in further diversification during adaptive radiation. Herein a polyploidy apart prospective gains also has many restrictions on fish biology that can dismiss the expected benefits (Gregory & Mable, 2005; Van de Peer *et al*., 2017). Nevertheless, the most researchers agreed that polyploid taxa are characterized by increased ecological and phenotypic plasticity that (i) facilitate to overcome the stressful and harsh environment and (ii) can promote to further diversification and speciation that is within ‘flexible stem’ hypothesis (West-Eberhard, 2003; Wund *et al*., 2008; Li & Guo, 2020; Van de Peer *et al*., 2021). Some polyploid lineages, for instance – Salmonidae and Coregonidae – represent nice examples of phenotypic and ecological plasticity along with possibility to evolve small-scale adaptive radiation especially within genera *Salvelinus*, *Salmo*, and *Coregonus* (Reshetnikov, 1980; Bernatchez *et al*., 1999; Alekseyev *et al*., 2002; Markevich *et al*., 2018; Segherloo *et al*., 2021; Levin *et al*., 2022).

The cyprinid fishes (fam. Cyprinidae sensu Tan & Armbruster, 2018) being one of the most diverse families of teleostean fishes (> 1780 species - Fricke *et al*., 2023) are the champions in the number of the polyplod species (ca. 600 species – Yang *et al*., 2022). Some of cyprinid polyploid lineages display bright examples of adaptive radiation based on trophic polymorphism (Nagelkerke *et al*., 1994; Roberts & Khaironizam, 2008; Levin *et al*., 2019; 2021; Qiao *et al*., 2020; Komarova et al., 2021). Among these, the most distinguished one in terms of the morpho-ecological diversification is the cyprinid genus *Labeobarbus* representing species-rich African hexaploid lineage (2n=150 – Oellerman & Skelton, 1990; Golubtsov & Krysanov, 1993) that comprises >130 species (Vreven *et al*., 2016; Fricke *et al*., 2023) and displays numerous trophically-based adaptive radiations (Nagelkerke *et al*., 1994; 2015; Mina *et al*., 1996; Sibbing & Nagelkerke, 2000; Dimmick *et al*., 2001; Golubtsov, 2010; Shkil *et al*., 2015; Levin *et al*., 2020). The *Labeobarbus* is widespread in Africa and presents in each of the ten African ichthyofaunal provinces (Vreven *et al*., 2016). Based on the recent data on phenotypic, ecological, and genetic data, the taxonomic diversity of the *Labeobarbus* might be greatly underestimated (Levin *et al*., 2019, 2020; Decru *et al*., 2022).

The Ethiopian Highlands is remarkably distinguished as a hotspot for multiple adaptive radiations of the *Labeobarbus* discovered during last 30 years (see Golubtsov *et al*., 2021). The most famous example of rapid adaptive radiation among non-cichlid fishes belongs to the lacustrine radiation of the *Labeobarbus* in Lake Tana, Ethiopia, where up to 15 species/ecomorphs were described that partitioned trophic resources (Nagelkerke *et al*., 1994; Mina *et al*., 1996; Zworykin *et al*., 2006; Shkil *et al*., 2015). Apart from the lacustrine radiation strongly predominating among fishes (e.g. Seehausen & Wagner, 2014), there are striking examples of the riverine adaptive radiations in non-cyprinid (Turner *et al*., 1985; Piálek *et al*., 2012; Esin *et al*., 2021; Říčan *et al*., 2021; Burress *et al*., 2022) as well as cyprinid fishes - *Labeobarbus*, in particular (Levin *et al*., 2019; 2020).

The most conspicuous phenotypic difference between sympatric ecomorphs of the *Labeobarbus* refers to mouth type variation. The core of mouth polymorphism is composed of four main phenotypes (Figure 1) that were recorded throughout Africa (Worthington, 1929; 1933; Matthes, 1963; Banister, 1973; Skelton *et al*., 1991; Nagelkerke *et al*., 1994): i) generalized (typical for barbs), ii) thick-lipped (‘rubberlip’), iii) scraping (‘chiselmouth’), and iv) large-mouthed. Mouth phenotypes are subjected to further sub-diversification and can include subtypes or variants that are recruited for occupying the new ecological niches (Levin *et al*., 2021b). Moreover, there are intermediate phenotypes possibly of hybrid origin (Banister, 1972; Vreven *et al*., 2019) that does not exclude their further specialization.

**Figure 1.**
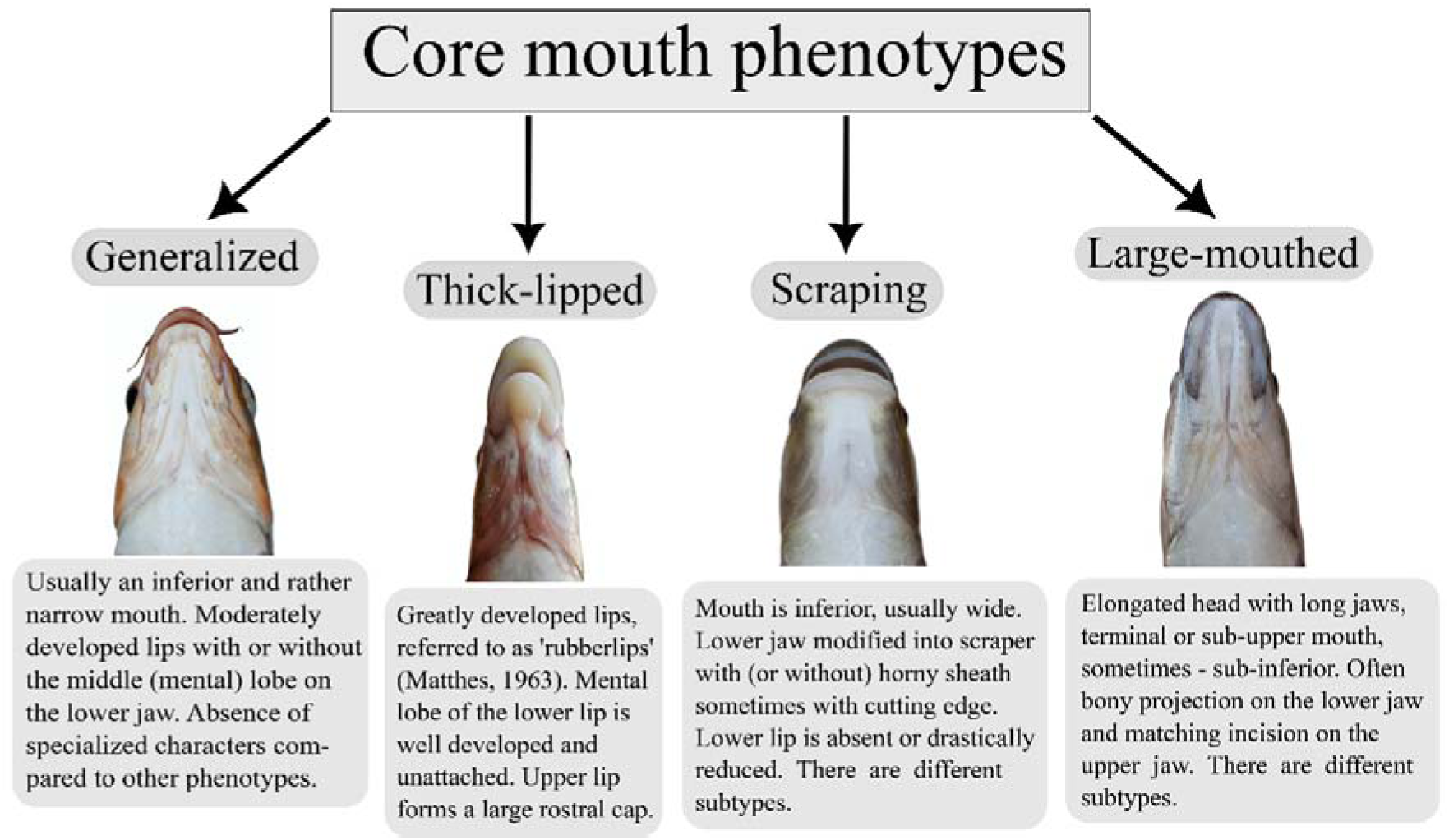
Pictorial scheme of core mouth phenotypes of the *Labeobarbus*. Photograph examples referred to the *Labeobarbus* from the rivers of the Ethiopian Highlands.

This outstanding mouth polymorphism of the *Labeobarbus* likely stems from ancestral inherited genetic templates since both parental lineages involved in the allopolyploidization event (Yang *et al*., 2015; 2022) belong to extant lineages and represent different mouth types. The contemporary representatives of the maternal lineage are characterized by generalized and thick-lipped mouth polymorphism (tetraploid Torinae, 2n = 100 - Roberts & Khaironizam, 2008; Walton *et al*., 2017; Coad, 2021) while paternal diploid lineage (*Cyprinion*, 2 n = 50, Ünlü *et al*., 1997; Esmaeili & Piravar, 2006) is represented by scraping mouth phenotype (Coad, 2021). Hence, mouth polymorphism of the *Labeobarbus* is ancestrally heritable, likely preadaptive and might re-evolve.

The parallel diversifications of the *Labeobarbus* in the riverine environment of the Ethiopian Highlands were recently discovered within the four geographically isolated drainages - i) the Didessa River in Blue Nile basin; ii) the Sore River in White Nile basin; iii) the Gojeb River in Omo-Turkana basin; and iv) the Genale River in Juba-Wabe-Shebelle basin, Indian Ocean catchment (Mina *et al*., 1998; Golubtsov, 2010; Levin *et al*., 2019; 2020; Golubtsov *et al*., 2021). Each diversification includes from four to eight sympatric ecomorphs divergent in mouth phenotype (Table 1) that presumably involves trophic adaptations. The genetic studies showed that all four riverine radiations of *Labeobarbus* were independently evolved from different ancestors although they are very closely-related based on mtDNA data that suggests recent and rapid diversification (Levin *et al*., 2013; 2019; 2020). Moreover, existing data suggests these radiations might be at various stages of the diversification (Levin *et al*., 2020; 2023 - in press). Of special interest is that these radiations could repeatedly emerge in the riverine environment that has been considered to be inappropriate for adaptive radiation due to heterogeneous and unpredictable conditions for a long time (reviewed in Burress *et al*., 2023).

**Table 1.**
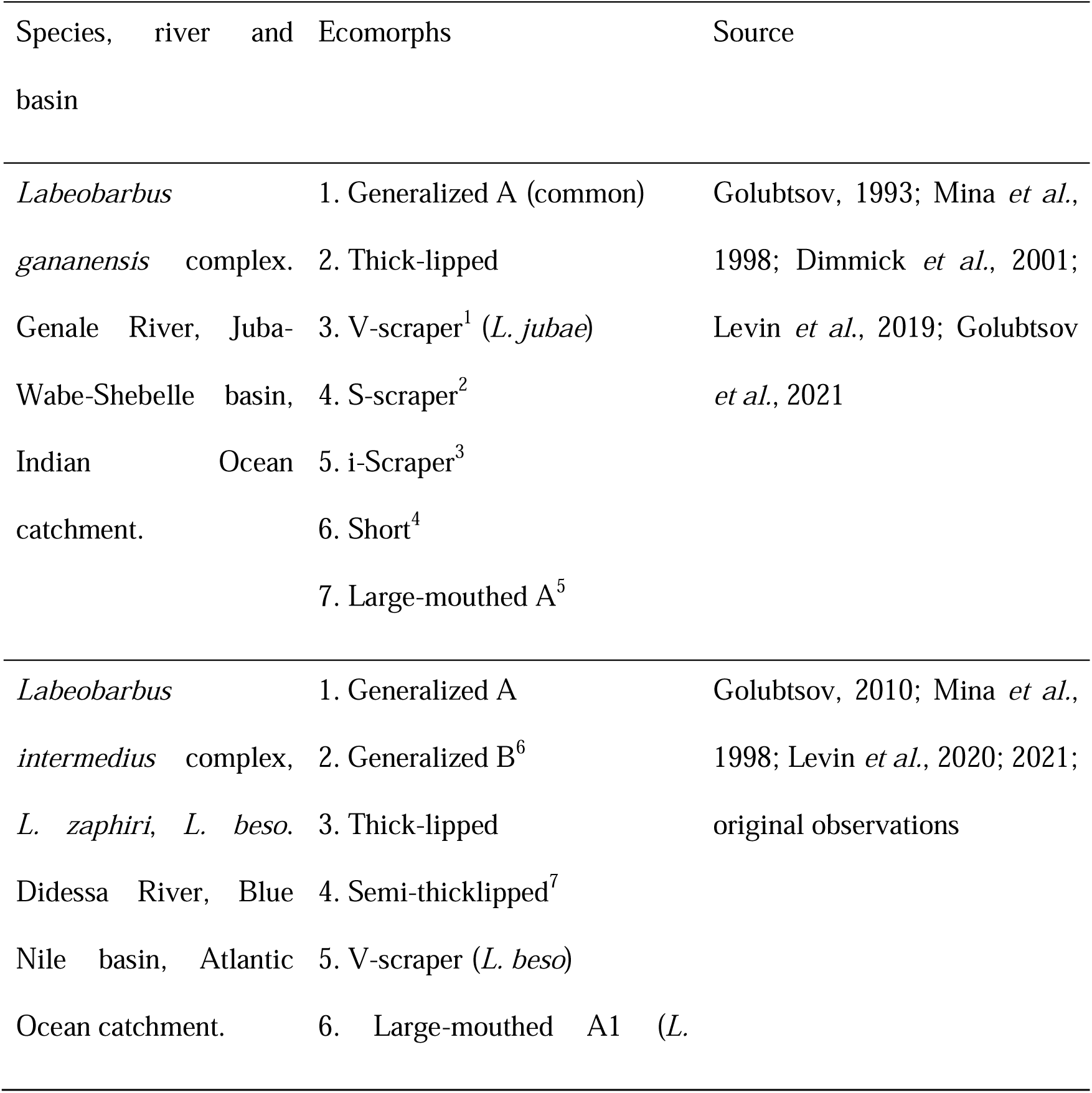
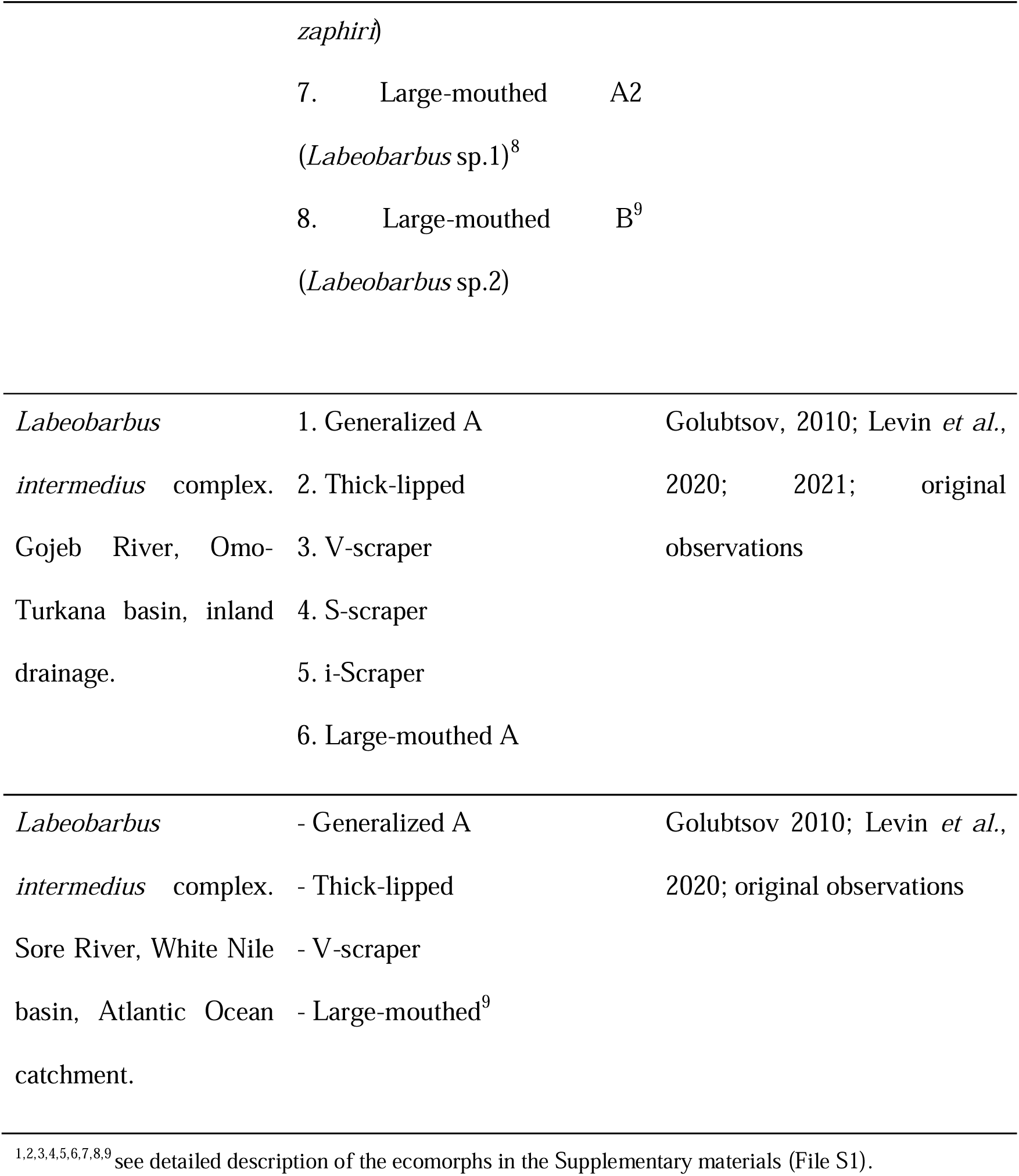
The composition of riverine sympatric ecomorphs of *Labeobarbus* spp. in isolated riverine basins in the Ethiopian Highlands (summarized from Golubtsov, 2010; Levin *et al*., 2019; 2020; 2021; Mina *et al*., 1998 and our unpublished data). The photographs of the fish ecomorphs are given in Figure 2.

A few studies on trophic ecology showed that sympatric riverine ecomorphs of the *Labeobarbus* bearing different mouth phenotypes have partitioned trophic resources - in particular, in the Genale River (Levin *et al*., 2019). Other riverine radiations of the *Labeobarbus* were not studied comprehensively from the point of trophic divergence. The recent but independent origin of similar diversifications in mouth phenotypes raises several questions. Do all diversifications partition trophic resources according to homologous mouth phenotypes? What is trophic specialization (if present) of various sympatric ecomorphs? Whether riverine diversifications comprising different numbers of ecomorphs / mouth (sub)phenotypes are at various stages of the process? If so, could patterns of the trophic diversification be sorted from more primitive to advanced aiming reconstruction of the trophic radiation? We try to answer these questions in a study of trophic diversification of sympatric ecomorphs in each of four *Labeobarbus* riverine radiations using the stable isotope composition and gut content.

## 2. Material and Methods

### Ethical approval

All applicable international, national, and institutional guidelines for the care and use of animals were strictly followed. All animal sample collection protocols complied with the current laws of Russian Federation and Federal Democratic Republic of Ethiopia.

### Study sites and material studied

Samples were collected under the framework of the Joint Ethiopian[Russian Biological Expedition (JERBE) in four rivers draining the Ethiopian Highlands and belonging to four major river basins (Fig. 2): (1) the Juba-Wabe-Shebelle basin in the Indian Ocean catchment – the Genale R. – 5.7025° N 39.5446° E; (2) the Blue Nile basin – the Didessa R., a tributary of the Blue Nile – 8.6921° N 36.4144° E; (3) the Omo-Turkana enclosed basin – the Gojeb R., a tributary of the Omo R. – 7.2539° N 36.7943° E; (4) the White Nile basin – the Sore R. ∼35 km downstream City Metu – 8.3987° N, 35.4378° E. Fish were caught by gill and cast nets in February–March 2011 (Didessa), March–April 2009, 2019 (Genale), February 2011 (Gojeb), April 2014 (Sore). Fish were killed with an overdose of MS-222 anaesthetic and then photographed using a Canon EOS 50D camera (Canon Inc. Tokyo, Japan).

**Figure 2.**
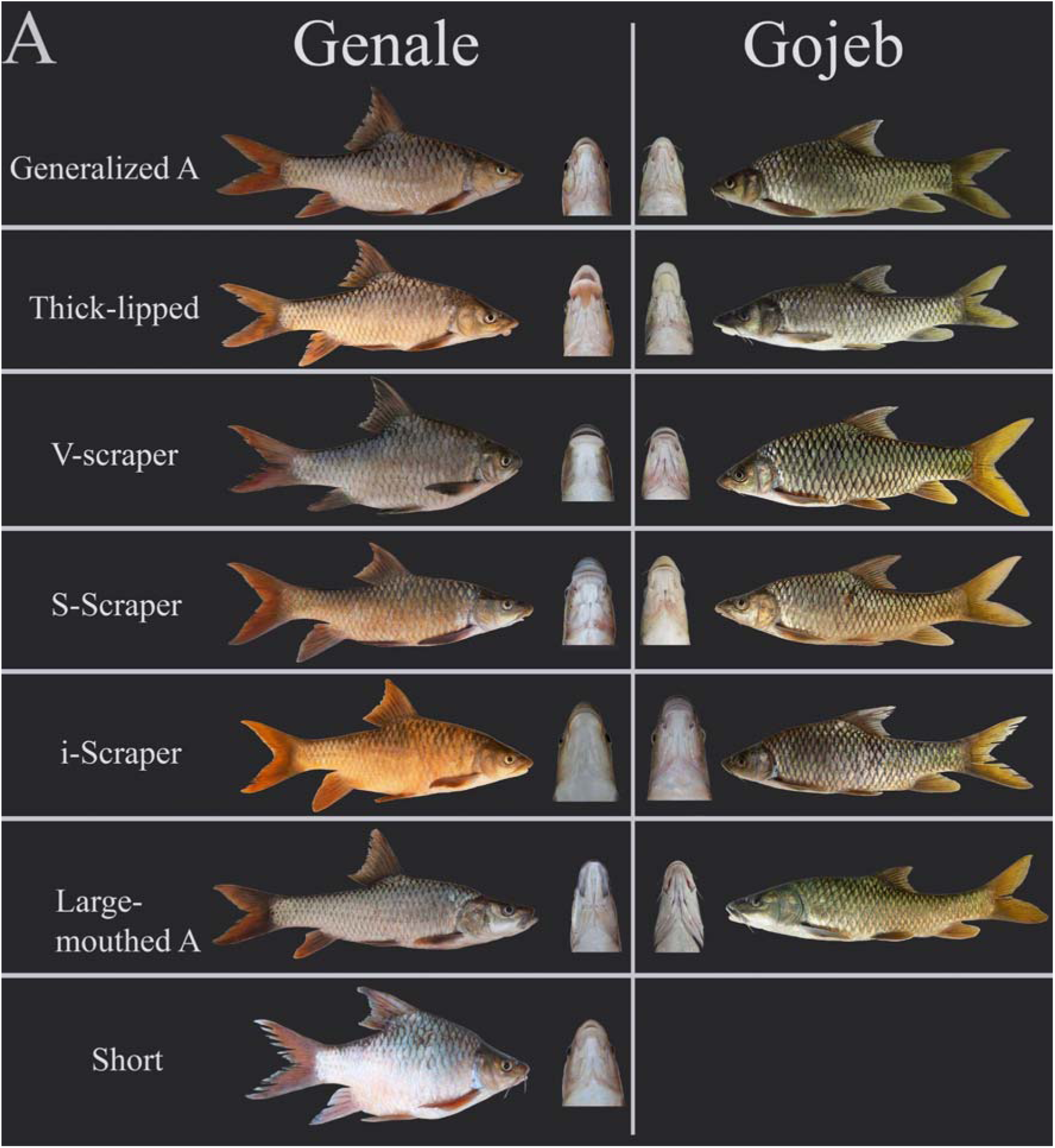
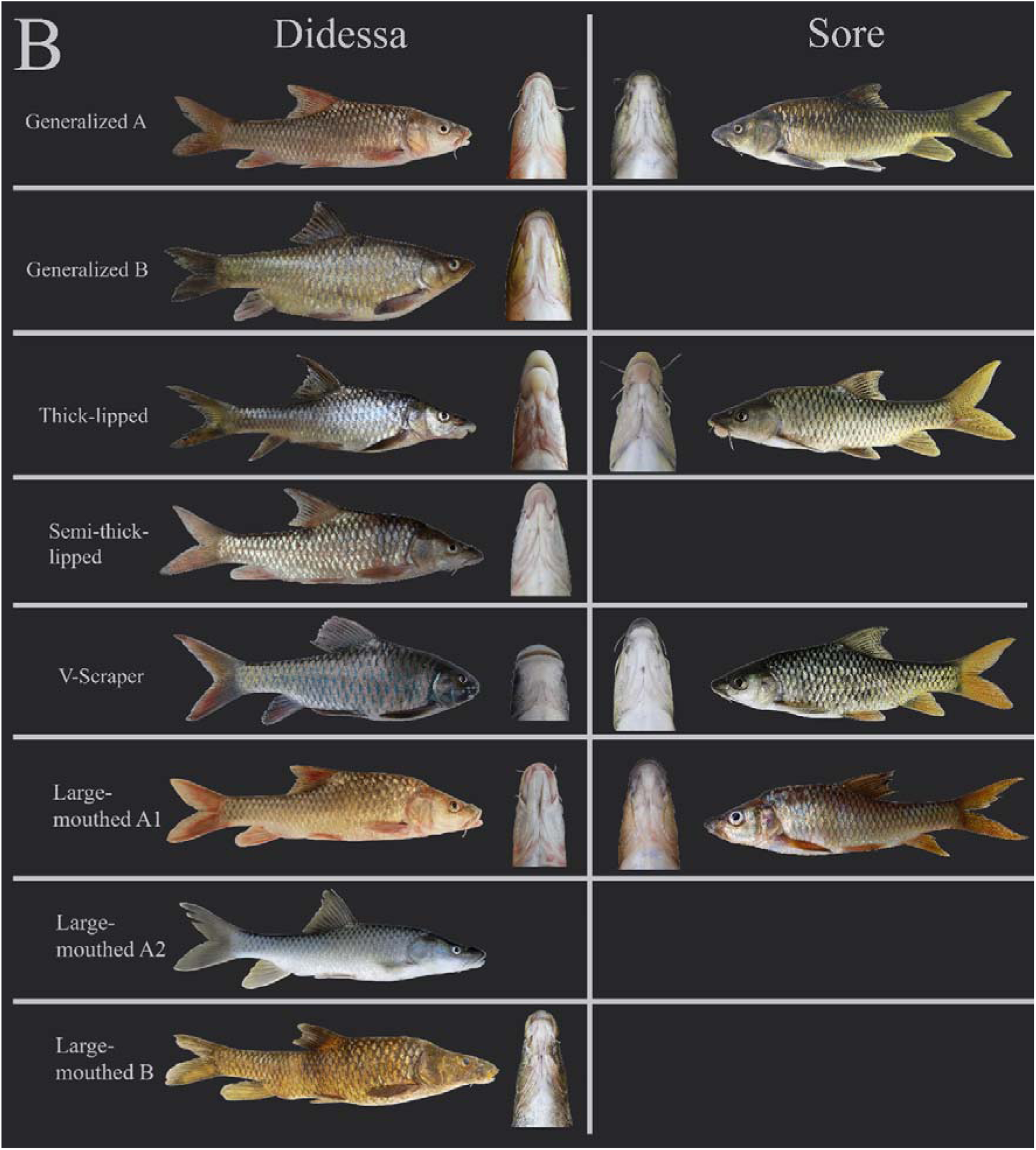
Appearance and mouth phenotypes of the ecomorphs of the *Labeobarbus* in different rivers of the Ethiopian Highlands: (A) the Genale and Gojeb Rivers; (B) the Didessa and Sore Rivers. An exceptionally rare representative of the Large-mouthed A2 ecomorph from the Didessa River is presented only from the lateral side.

**Figure 3.**
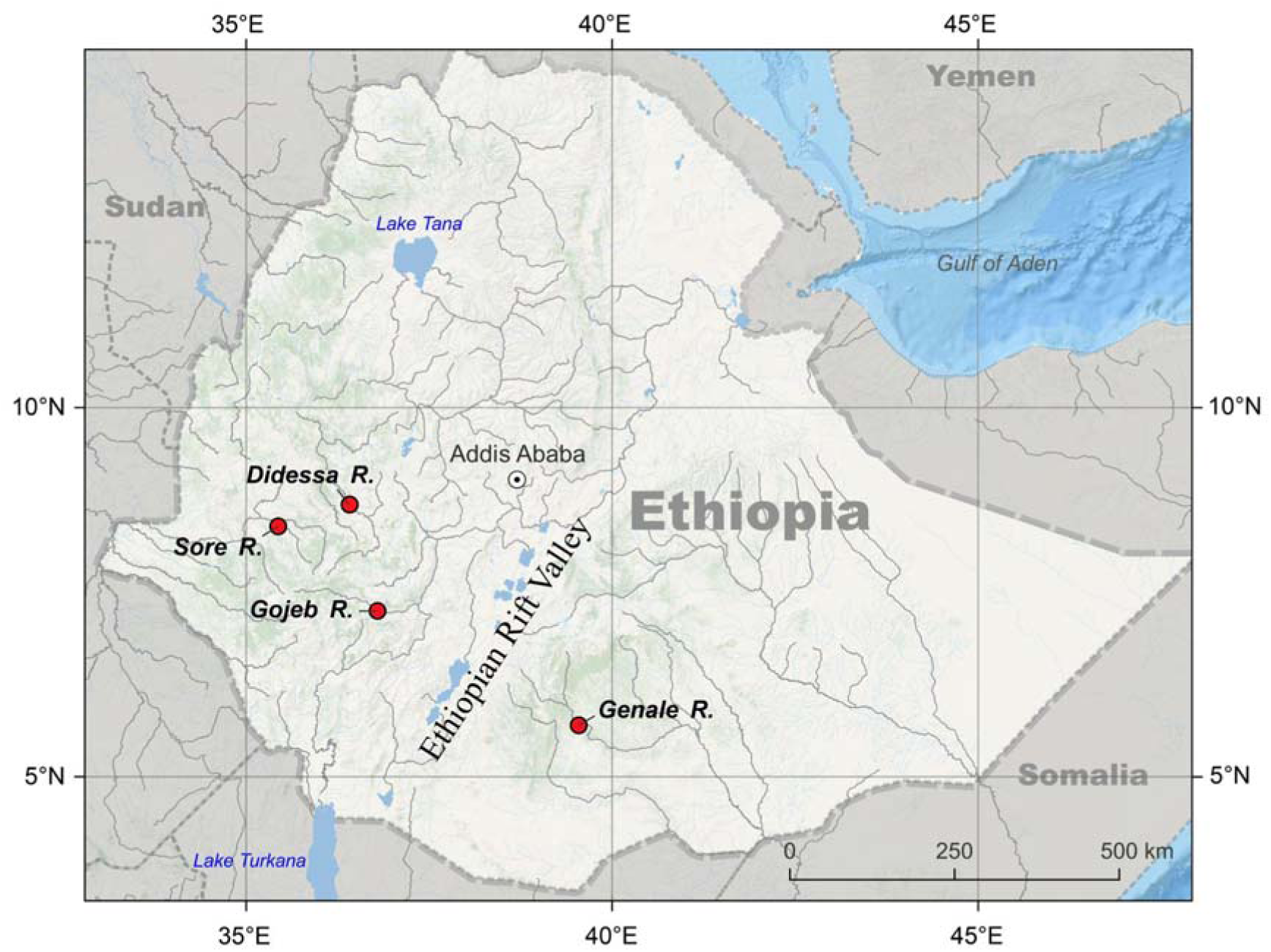
Map of sampling sites of the four riverine radiations of the Ethiopian *Labeobarbus*.

Standard length (SL, mm) and gut length (GL, mm) were measured with a ruler. Fish were preserved first in 10% formalin and then transferred to 70 % ethanol. Gut length was studied in 461 individuals. All specimens were deposited at A.N. Severtsov Institute of Ecology and Evolution, the Russian Academy of Sciences, Moscow under provisional labels of JERBE. The detailed information on material studied (gut length, stable isotope composition and gut content) is given in Supporting Table S2.

### Stable isotopes composition

For stable isotope (SI) analyses, white muscle tissue from the dorsal side of the body under the dorsal fin was sampled from freshly collected specimens. White muscle samples were dried at 60 °C. The samples were weighed using a Mettler Toledo MX5 microbalance (Mettler Toledo, Columbus, OH, United States) with 2 μg accuracy, and wrapped in tin capsules. The weight of the fish tissue samples varied from 250 to 500 μg. SI analysis was conducted at the Joint Usage Center of the A.N. Severtsov Institute of Ecology and Evolution RAS, Moscow. Briefly, a Thermo Delta V Plus continuous-flow IRMS was coupled to an elemental analyzer (Flash 1112) equipped with a Thermo No-Blank device. The isotopic composition of N and C was expressed in the δ notation relative to the international standards (atmospheric nitrogen and VPDB, respectively): δX (‰) = [(Rsample/Rstandard) − 1] × 1000, where R is the molar ratio of the heavier and lighter isotopes. The samples were analyzed with a reference gas calibrated against the International Atomic Energy Agency (IAEA) reference materials USGS 40 and USGS 41 (glutamic acid). The measurement accuracy was ± 0.2 ‰. Along with the isotopic analysis, the nitrogen and carbon content (as %) and C/N ratios were determined. In total, 400 white-muscle samples were analyzed.

### Diet analysis

Following the best practice for estimation of trophic niches the in tropic rivers (e.g., Davis *et al*., 2012) we studied gut content in addition to SI composition. Gut content was extracted from preserved specimens, dried on filter paper and weighed using a Pioneer PX84/E balance with 0.0001 g accuracy. The diet particles were identified using Olympus CX41 microscope (100–1000× magnification) and Motic DMW-143-N2GG stereomicroscope (100–400× magnification). The diet components were grouped into: (i) detritus, (ii) invertebrates, (iii) macrophytes, (iv) periphyton, (v) filamentous algae, (vi) fish (body remnants and scales), and (vii) mineral ground. The group ‘Invertebrates’ included mainly the larvae and imago of amphibiotic insects and their fragments as well as imago of aerial insects (Coleoptera, Hymenoptera); rarely Cladocera. The group ‘Macrophytes’ included any fragments of helophytic and semi-aquatic plants - such as leaves, stems or seeds. A composite measure of diet, an index of relative importance (IR) [Natarajan & Jhingran, 1961; Popova & Reshetnikov, 2011], was used to assess the contribution of different components to the diet. The IR index was calculated as follows: IR = (Fi × Pi)/(∑(Fi × Pi)) × 100%, where Fi = the frequency of occurrence of each food group, and *Pi* = its part by weight; the value of *i* itself changes from 1 to n (n = the part of food organisms in the food bolus). In total, 195 food boluses were analyzed for gut content (Supplementary Table S2).

### Statistical analyses

Several R packages and functions were used for the statistical analyses and plot construction. Basal descriptive statistics was obtained using the *summarytools* library [Comtois, 2022]. Central tendencies in the text are presented as means and 1SD. The Kruskal-Wallis test was applied for comparisons of the values of gut length GL and SI composition using the function *kruskalTest* in each radiation (in library *PMCMRplus*) with subsequent pairwise comparisons using post-hoc Dunn’s all-pairs test with Bonferoni adjustment - function *kwAllPairsDunnTest* [R Core Team, 2023]. The violin boxplots were obtained using the *ggplot2* library [Wickham, 2022]. The package SIBER v.2.1.6 [Jackson *et al*., 2011] was used to assess the differences in the isotopic trophic niche features. The total convex hull areas (TA), core trophic niche breadths, and sample size-corrected standard ellipse area (SEAc) were estimated. The trophic overlap for 95% TA was estimated using nicheROVER [Lysy et al., 2021], a method that is insensitive to the sample size and incorporates statistical uncertainty using Bayesian approach [Swanson *et al*., 2015].

## 3. Results

### 3.1. Diet

Food spectra of *Labeobarbus* spp. were rather diverse. Feeding of sympatric ecomorphs in all four rivers was divergent but at various degrees as estimated by index of relative importance (Figure 4). Detailed description of the diet for each ecomorph is given in Supplementary material S3.

**Figure 4.**
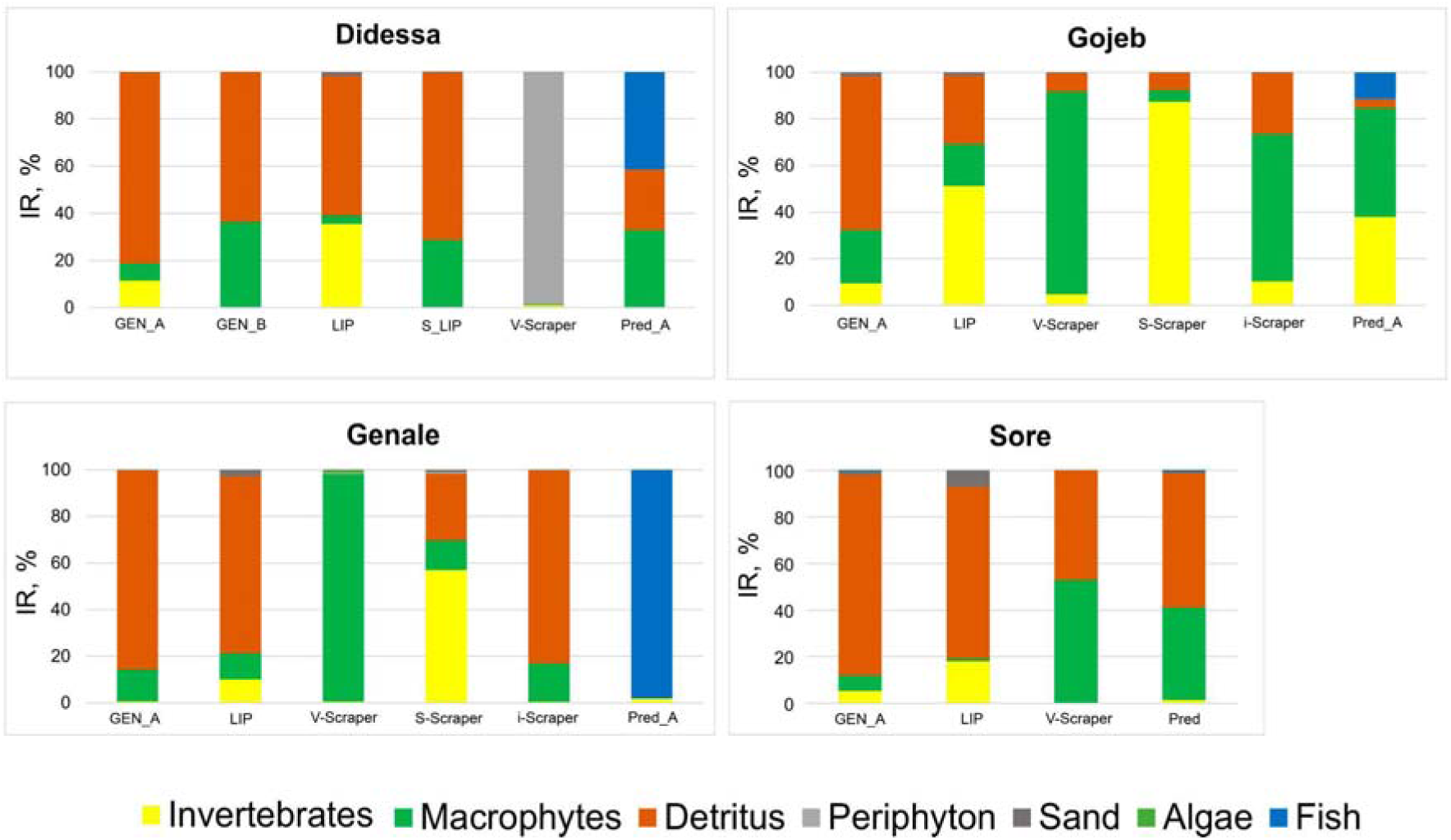
Food spectra (IR: the index of relative importance) of the sympatric ecomorphs of the *Labeobarbus* spp. from the Didessa, Gojeb, Genale, and Sore rivers. Abbreviations of the ecomorphs: GEN_A - Generalized A; GEN_B - Generalized B; LIP - Thick-lipped; S_LIP - Semi-thicklipped; Pred_A - Large-mouthed A; Pred_B - Large-mouthed B.

Based on the food spectra, up to five trophic specializations can be detected in the Didessa River: i) detritivore (Generalized A), ii) detritivore-macrophytophage (Generalized B and Semi-thicklipped), iii) detritivore-invertivore (Lipped), iv) periphyton feeder (V-scraper), and v) piscivore-omnivore (Large-mouthed A and B). Notably, detritus seems to be a core food for many ecomorphs except for the V-Scraper and Large-mouthed. A sub-specialization within the piscivory strategy is possible taking into account the drastic difference in mouth structure (Figure 2B) and diet between the Large-mouthed ecomorphs. Seemingly, up to five feeding strategies can be distinguished in the Gojeb River: i) detritivore (Generalized A), ii) invertivore-detritivore (Lipped); iii) macrophytophagous (V-Scraper and i-Scraper), iv) invertivore (S-scraper), and v) omnivore-piscivore (Large-mouthed A). Compared to the Didessa, a detritus was a core food only for the Generalized A ecomorph. Large-mouthed ecomorph is not predominantly piscivorous but rather omnivorous with a small portion of fish food. In the Genale River, up to four feeding strategies can be detected – i) detritivore (Generalized A, Lipped, and i-Scraper), ii) macrophytophagous (V-Scraper), iii) invertivore (S-scraper), and iv) piscivore (Large-mouthed A). Remarkably, a detritus was a core food for three ecomorphs in the Gojeb River, while the Large-mouthed ecomorph is a strongly piscivorous specialist. In contrast, little diversification in food spectra was detected in the Sore River. Briefly, two feeding strategies can be recognized: i) detritivore (Generalized A, Lipped, and i-Scraper), and ii) macrophytophagous-detritivore (V-Scraper and Large-mouthed). Remarkably detritus was a core food for three of four ecomorphs although a notable portion (almost 20 %) of benthic invertebrates was detected in the Lipped ecomorph. Large-mouthed ecomorph is not a piscivorous specialist in the Sore River.

### 3.2. Stable Isotope Composition

Basic statistics for both δ^15^N and δ^13^C values and data on the total area (TA), standard ellipse area (SEA), and corrected standard ellipse area (SEAc) are given in Supplementary Materials S4. Detailed results are presented in subsequent sections.

#### 3.2.1. Didessa Radiation

Among seven ecomorphs, several were significantly divergent in SI values (six pairwise comparisons in both δ^15^N and δ^13^C - Table S4). The largest values of δ^15^N were detected in two Large-mouthed (piscivory) ecomorphs (12.5±0.6 and 12.9‰) as well as in highly specialized V-scraper *L. beso* (12.3±0.6‰). The V-scraper was also most enriched in ^13^C (δ^13^C -17.7±2.4%) (Figure 5). The lowest values of both δ^15^N (8.33±0.7%) and δ^13^C (-22.8±0.9%) were in the Generalized B ecomorph. Other ecomorphs, Generalized A, Thick-lipped, and Semi-thicklipped, had intermediate δ^15^N and δ^13^C values, although Thick-lipped ecomorph had somewhat higher δ^15^N values (10.6±1.3‰) compared to both Generalized A and Semi-thicklipped (8.3±1.3 and 9.2±1.1‰, respectively). The maximum difference in mean δ^15^N values between ecomorphs in the Didessa was 4.61‰. The highest difference between ecomorphs in the mean δ^13^C values was 5.10‰. The isotopic niches (as assessed by the standard ellipses) were almost fully separated between Generalized B and V-Scraper (overlap <1%), between Generalized B and Large-mouthed A (overlap <1%), and weakly overlapped between Semi-Thicklipped and V-scraper, Semi-Thicklipped and Large-mouthed A, and Generalized A and V-Scraper (overlap <6%, <10 %, and <11%, respectively) (see details in Supplementary File S4). The most overlapping trophic niches were between Generalized A and B (57 and 90%) and between Generalized A and Semi-Thicklipped (77 and 79%) (Supplementary File S4).

**Figure. 5.**
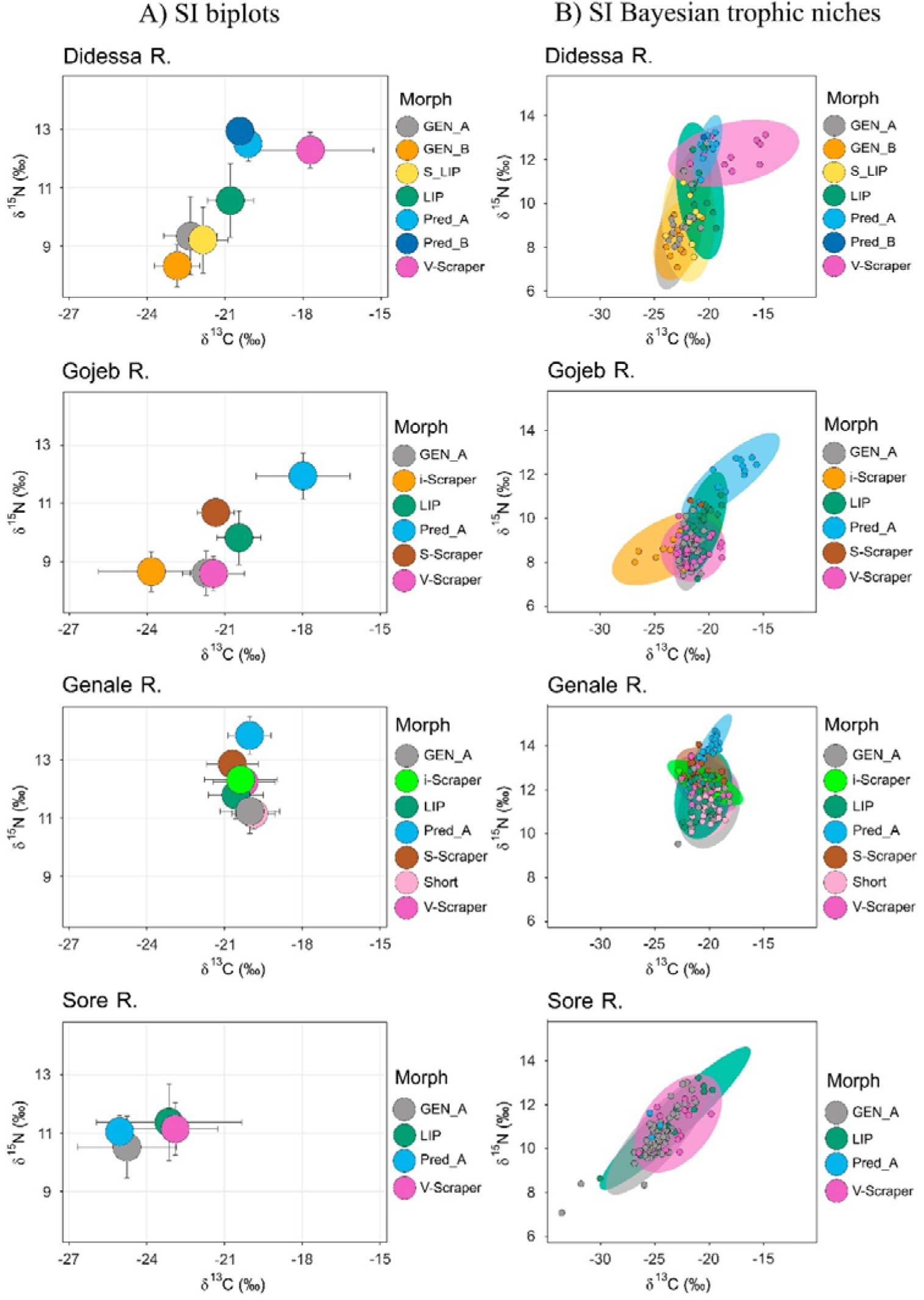
Stable isotope biplots (A) showing mean values and 1 SD, and Bayesian ellipses showing trophic niche widths and overlaps (B) in sympatric ecomorphs of the *Labeobarbus* spp. from the Didessa, Gojeb, Genale, and Sore Rivers. Ellipses with 95% confidence intervals are based on standard ellipses corrected for small sample sizes (SEAc; isotopic niche metrics; SIBER package). Each point corresponds to the individual isotopic value. Abbreviations of the ecomorphs: GEN_A - Generalized A; GEN_B - Generalized B; LIP - Thick-lipped; S_LIP - Semi-thicklipped; Pred_A - Large-mouthed A; Pred_B - Large-mouthed B.

#### 3.2.2. Gojeb Radiation

Among six ecomorphs, some were significantly divergent from others in SI values (five and six pairwise comparisons in δ^15^N and δ^13^C values, respectively; Table S4). The largest values of δ^15^N and δ^13^C were found in Large-mouthed (piscivory) ecomorph (11.94±0.8‰ and -17.98 ±1.8‰, respectively) that was notably (sometimes significantly - Figure 5, Table S4) higher than in all other sympatric ecomorphs. The minimal values of δ^15^N were detected in the V-scraper (8.60±0.6‰) and Generalized (8.61±0.8‰) ecomorphs while in the i-Scraper ecomorph the δ^13^C value was at the minimum (-23.84±2.0 ‰). The total divergence between ecomorphs was lower in δ^15^N but higher in δ^13^C values compared to the Didessa radiation. In particular, the highest difference between ecomorphs in the mean δ^15^N values achieved 3.34‰, while the same for the mean δ^13^C values was 5.86‰ (Figure 5, Table S4). The isotopic niches were almost fully separated between Large-mouthed and V-Scraper (overlap <2%) and between Generalized and Large-mouthed (<4%) while the most overlapping trophic niches were between Generalized and V-Scraper (overlap 70 and91%) (Supplementary File S4).

#### 3.2.3. Genale Radiation

Among seven ecomorphs, some were significantly divergent from others in δ^15^N values (eight pairwise comparisons), but not in δ^13^C values (Table S4). The largest value of δ^15^N was found in the Large-mouthed (piscivory) ecomorph (13.83±0.7 ‰) (Figure 5, Table S4). The minimal values of δ^15^N were detected in the Short and Generalized ecomorphs (11.15±0.5‰ and 11.23±0.8‰, respectively). The range between largest and lowest mean δ^13^C values was small, from -19.87±0.8‰ (Short) to -20.69±1.0‰ (S-Scraper). The total divergence between ecomorphs was lower in δ^15^N values than in both the Didessa and Gojeb radiations. In particular, the highest difference between ecomorphs in mean δ^15^N values was 2.73‰, and only 0.82‰ in δ^13^C values. In the isotopic niches the most divergent ecomorph was the Large-Mouthed, which overlap with other ecomorphs ranged from 0.06% (vs. Short) to 38 and 43% (vs. benthophagous S-Scraper). Among other comparisons a low overlap was detected between S-Scraper and Short (7 and 8%) (Supplementary File S4).

#### 3.2.4. Sore Radiation

Among four ecomorphs, a few were significantly divergent in δ^13^C values only (two pairwise comparisons - Generalized vs. Thick-Lipped and Generalized vs. V-Scraper; Table S4). The highest difference between ecomorphs in δ^15^N values was 0.85‰ only, while in the mean δ^13^C values it was 1.86‰. The isotopic niches greatly overlapped (67…82%) in all comparisons (Supplementary File S4).

### 3.3. Gut Length

The relative gut length varied significantly within each radiation except for the Sore River (Figure 6). In three of four radiations, there were significant differences between some ecomorphs (Figure 6; Supplementary File S5). Longest guts (up to 548-692% SL) were detected among V-scrapers (in the Didessa, Genale, and Gojeb Rivers). The large-mouthed ecomorphs showed shortest guts (highest values were up 194-228% SL) except for the Sore River, where it reached 251±75%. Other ecomorphs had intermediate gut length; sometimes they differed from each other like Generalized and Thick-lipped ecomorphs in the Gojeb River (Figure 6). In spite of diverse mouth phenotypes in the Sore River, the ecomorphs were not divergent from each other representing the middle length of the gut (means varied within 251-290% SL).

**Figure 6.**
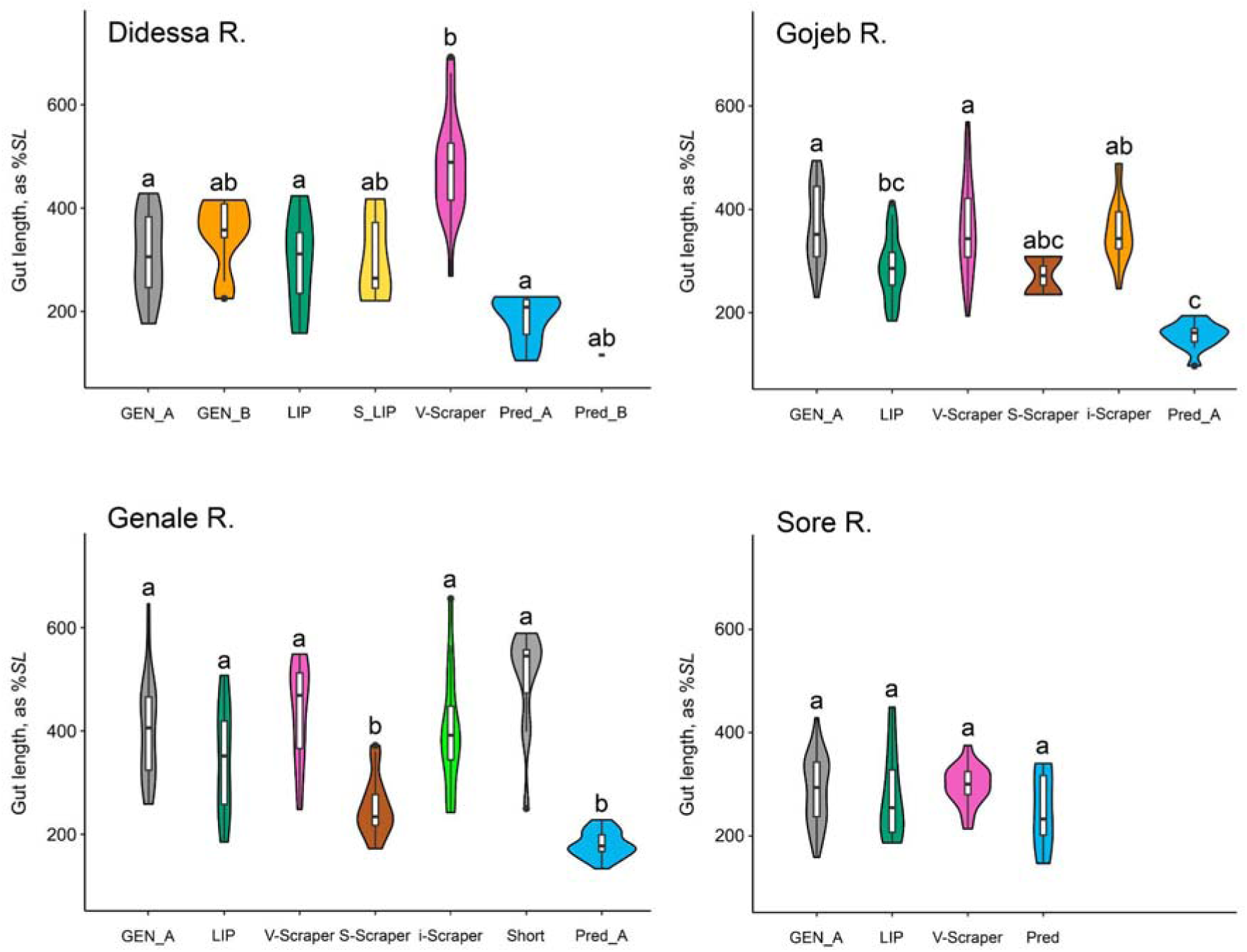
Violin plots of relative gut length distribution in sympatric ecomorphs from the four riverine radiations of the *Labeobarbus*. Min-max values (whiskers), 1st and 3rd quartiles (white vertical bars), median values (black horizontal bars), and outliers (black points) are indicated. Letters above the violin plots indicate significant differences between ecomorphs (p < 0.05, Kruskal–Wallis test with Dunn’s post hoc test). Abbreviations of the ecomorphs: GEN_A - Generalized A; GEN_B - Generalized B; LIP - Thick-lipped; S_LIP - Semi-thicklipped; Pred_A - Large-mouthed A; Pred_B - Large-mouthed B.

## 4. Discussion

Our results show that parallel trophic diversifications occurred within the polyploid lineage, the genus *Labeobarbus*. These diversifications differ in degrees of specialization within the same set of core mouth phenotypes (generalized, thick-lipped, scraping, and large-mouthed), indicating various stages of the process. We discuss the results obtained in the context of trophic specializations of certain mouth phenotypes, the preadaptive nature of the ancestral discrete mouth polymorphism, which likely originated via allopolyploidzation of the *Labeobarbus* lineage, and attempt to reconstruct the process of trophic diversification by analyzing four repeated cases discovered in the riverine environment of the Ethiopian Highlands.

### 4.1. Trophic resource partitioning and trophic specializations

Our study revealed that differences in diet between ecomorphs of the *Labeobarbus* in each river were generally confirmed by SI analysis. For instance, the large-mouthed ecomorphs which had piscivorous diet were also significantly enriched in δ^15^N values (Figures 4 and 5). In contrast, the ecomorphs with detritivorous mode of feeding had lowest δ^15^N values. The periphython feeder that have been recorded in the Didessa River had a high level of δ^15^N values comparable with such in piscivorous ecomorphs but was characterized by enriched δ^13^C values (Figure 5). The invertivorous ecomorphs whose diet was based on the larvae of amphibiotic insects had intermediate levels of δ^15^N values between detritivores and piscivores. The data on gut length were less informative although the piscivorous ecomorphs had significantly shortened gut (Figure 6) that is in line with literature data for both *Labeobarbus* (Nagelkerke et al., 1994; Levin et al., 2019) and other fishes (e.g., Hugueny & Pouilly, 1999; Wagner et al., 2009). We summarized obtained data on the trophic specialization of sympatric ecomorphs from all rivers based on the diet, gut length, and SI composition in Table 2.

**Table 2.**
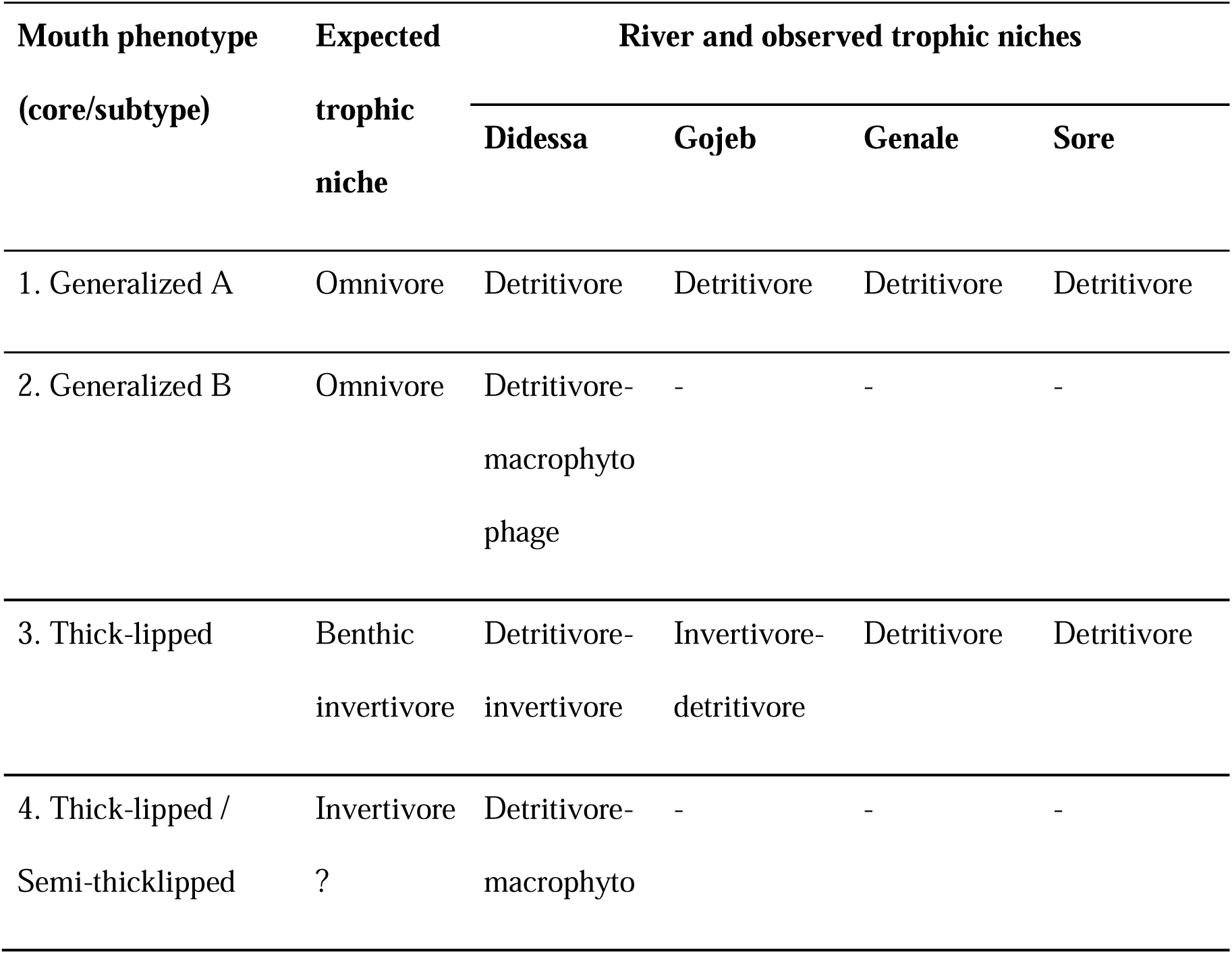
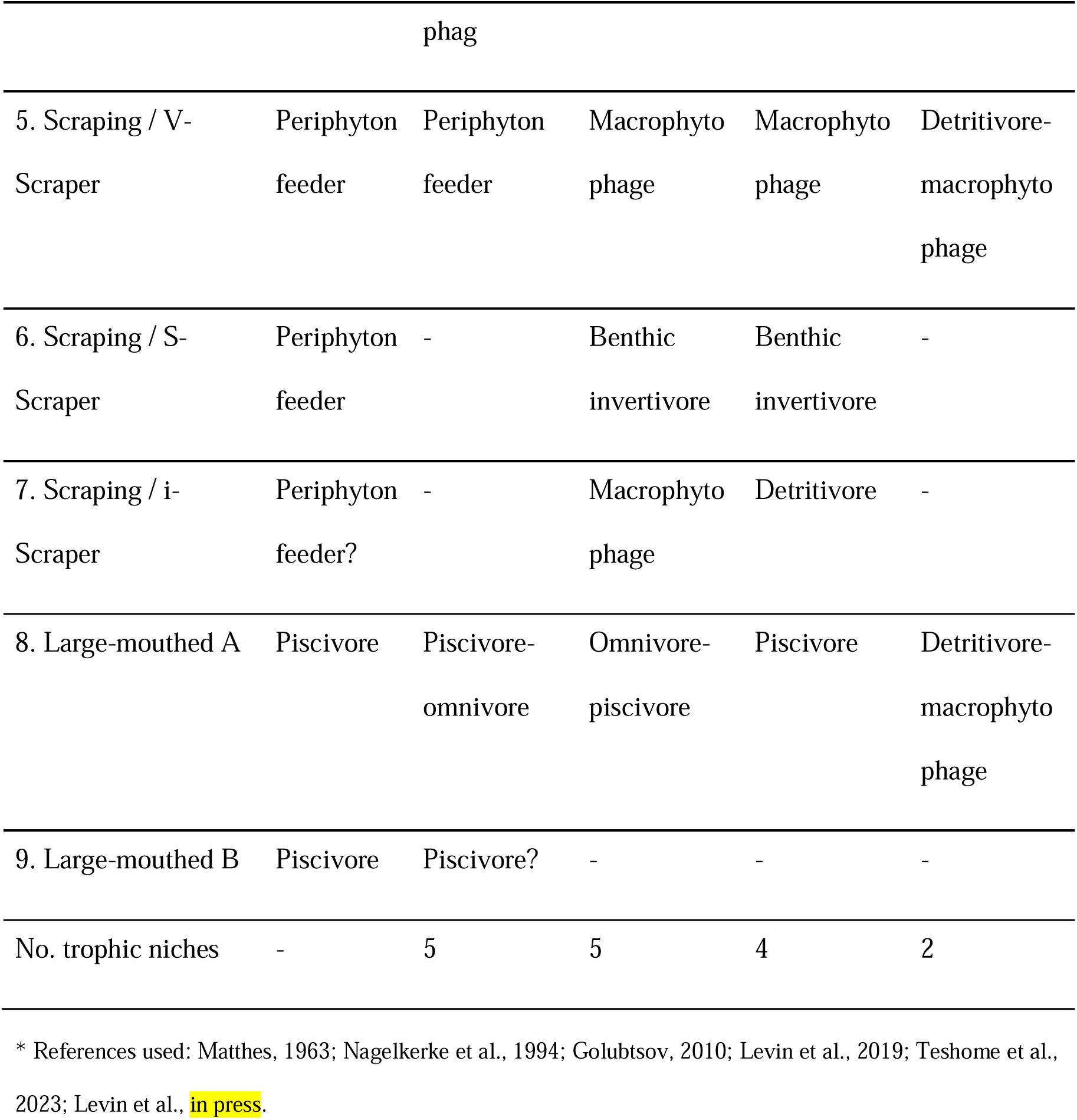
Correlation between mouth phenotype and trophic specialization in the *Labeobarbus* spp. from four riverine radiations. The expected trophic niche is based on prediction from the mouth phenotype and published data on trophic ecology*. Observed trophic niche - data obtained in this study.

Results of this study suggest five main trophic strategies among sympatric ecomorphs of the *Labeobarbus*: i) detritivory, ii) macrophytophagy, iii) invertivorous benthophagy, iv) periphyton feeding, and v) piscivory. Apart those, some ecomorphs had mixed modes of feeding - e.g., detritivory-macrophytophagy, detritivory-invertivory, and piscivory-omnivory (also with inclusion of a noted portion of detritus). Their more flexible trophic specialization might be an adaptive strategy for living in mountain rivers with unstable hydrological regimes (Jepsen & Winemiller, 2002). Although the core of mouth phenotypes (generalized, thick-lipped, scraping, and large-mouthed) was the same in four riverine radiations of the *Labeobarbus*, some phenotypes display more diversified sets of ecologically relevant mouth subtypes. Scraping mouth phenotype was represented by three subtypes, which occupy four different trophic niches (periphyton feeding, detritophagy, invertivorous benthophagy, and macrophytophagy). It is an outstanding example of trophic diversification of scraping mouth phenotype that previously was considered as adapted for feeding via scraping the algal periphyton from stones and rocks (Matthes, 1963; see also Vreven et al., 2016). Recent studies uncovered similar patterns of diversifications within other phylogenetically distant cyprinid lineages bearing the scraping mouth phenotype - the *Garra* and *Schyzopygopsis*, in particular (Komarova et al., 2021; 2022; Levin et al., 2021a). Contrary to the *Labeobarbus* radiations that likely initiated from the generalized ancestor as most ubiquitous throughout the generic range, the highly specialized scraper lineages (*Garra* and *Schyzopygopsis*) could give a trophic radiation outside the ancestral narrow specialization.

The Large-mouthed phenotype of the *Labeobarbus* demonstrated further sub-specialization within the adaptive zone of the piscivory in the Didessa River while in other rivers this phenotype had only one feeding strategy that varied from obligate piscivory (Genale) via piscivory-omnivory (Gojeb) to non-piscivory mode of feeding (Sore). The sub-specialization of the Large-mouthed phenotype in the riverine environment is similar in some extent to the more diversified set of specializations discovered in Lake Tana where up to seven sympatric large-mouthed species/ecomorphs were described (Nagelkerke et al., 1994), whose piscivory specialization subdivided in benthic and pelagic zones (de Graaf et al., 2010).

A large incongruence between mouth phenotype and expected trophic niche is noted not only for the trophic specialists. The Generalized ecomorph, including Ethiopian species and populations, was considered omnivorous (Matthes, 1963; Teshome et al., 2023), but according to our data it is detritivorous in all studied rivers (Levin et al., 2019; 2023 – in press; this study). At the same time further diversification within Generalized mouth phenotype (A and B subtypes) have been found. It is the emergence of an ecomorph with horseshoe shape of the lower jaw without mental lobe and deep body in the Didessa River (Generalized B – Figure 2B). This ecomorph had lowest both δ^15^N and δ^13^C values among other sympatric ecomorphs occupying a niche of detritivore-macrophytophage (Figures 4-5).

Noticeably, a detritus was the main food not only for generalists but often for thick-lipped phenotype and also for all ecomorphs from the Sore River regardless their mouth phenotypes. We consider detritus as the most available and permanent trophic resource in riverine ecosystems of the Ethiopian Highlands with an unstable hydrological regime. In fact, it is a core food for the Generalized ecomorph and an important food resource for other ecomorphs.

Apart from the core mouth phenotypes, the intermediate ones were also found. For instance, i-Scraper phenotype is intermediate between the Scraping and Generalized. Its hybrid origin in the Genale River (by S-scraper and Generalized) was confirmed by genetic data (Levin et al., 2019). Isotopic niche of i-Scraper from the Genale was between parental species (Figure 5) but the gut content was similar to sympatric Generalized ecomorph. Supposedly, i-Scraper intermediate phenotype might have its own trophic sub-niche. Another example of intermediate mouth phenotype that seemingly occupies its own sub-niche refers to the Semi-thicklipped ecomorph (see Figures 4-5) in the Didessa River. Obtained results are consistent with the syngameon hypothesis, proposing that hybridization between members of the radiation can promote further niche expansion and diversification (see Seehausen, 2004; Frei et al., 2022). Intermediate mouth phenotypes were reported among the *Labeobarbus* – between scraping ‘*Varicorhinus*’-like and generalized phenotypes by Banister (1972) and Nagelkerke and Sibbing (1996) as well as between scraping and thick-lipped mouth phenotypes by Vreven et al. (2019). In the last case the intermediate phenotype resembled the generalized one (Generalized B in our study). Authors revealed a hybrid nature of this phenotype by genetics (Vreven et al., 2019) but there was no data on trophic ecology or specialization. Actually, the recruiting of novel trophic niches via hybridization is an intriguing issue that is yet weakly studied (but see experimental study of Selz and Seehausen, 2019).

In summary, the *Labeobarbus* trophic radiations involve a mixture of pelagic predator (piscivore) and a wide assortment of benthic-oriented specialists. The benthic-pelagic habitat axis is most frequent in cyprinid and cichlid adaptive radiations in both lakes and rivers (Nagelkerke et al., 1994; Cooper et al. 2010; Levin et al., 2019; Burress et al., 2023).

### 4.2. Decoupled form and function – preadaptive phenotypes and their functionalization

Our results show that the same mouth phenotypes in the *Labeobarbus* may occupy different trophic niches. A mismatch between expected and observed trophic specializations may have different nature: i) specialized phenotype has unspecialized (generalized or omnivorous) feeding mode that is known as Liem’s paradox (Liem, 1980; Robinson et al., 1998), and ii) specialized phenotype has unexpected (biased) trophic specialization possibly due to insufficient knowledge on trophic specializations in certain lineages. We focus in this study on the first phenomenon. Liem’s paradox was established for cichlid fishes (e.g., Liem, 1980; Sturmbauer et al., 1992; Wagner et al., 2009; Binning et al., 2009; Torres-Dowdall & Meyer, 2021) but it was also detected in other taxonomic groups (e.g., in cyprinids – Lammens et al., 1991; Levin et al., 2021a; Komarova et al., 2022). The mismatch between form and function might be i) a temporary phenomenon for proper specialist explained by plasticity of its diet in some circumstances (ecological release that may be a base for evolutionary re-specialization – see above-mentioned example of highly specialized scraping periphyton feeders of the genus *Garra* that could re-specialize in other trophic specializations – Komarova et al., 2022) or ii) heterochronous decouple of form and function when the emerging phenotype is not yet functionalized, i.e. phenotype is preadaptive (expectation on its trophic adaptation based on phenotypic features) but not yet involved in trophic resource partitioning. This is exactly the case of the mouth diversity in the Sore River, where detritus was a main food for all mouth phenotypes including the Large-mouthed, i.e. mouth divergence was uncorrelated with the use of food resources. In other words, mouth polymorphism in the Sore River is ecologically almost non-functional compared to other riverine diversifications. This contradicts the ‘ecological theory’ of adaptive radiation in neglecting the ‘habitat first rule’ in particular (see Schluter, 2000) but fits the flexible stem hypothesis (Wund et al., 2008; Gibert, 2017) or its variant known as the ‘plasticity-first’ evolution (Levis & Pfennig, 2016; 2019). According to this hypothesis, an adaptive phenotypic plasticity in an ancestral population could precede adaptation to a new environment through the process of genetic assimilation. Remarkably, the case of decoupled form and function in the *Labeobarbus* is shared to some extent with such in riverine adaptive radiations in the pike cichlids, genus *Crenicichla* (Burress et al., 2023) that may be a general feature for young radiations in riverine environments.

As already mentioned, the repeated mouth polymorphisms of the *Labeobarbus* may be predetermined due to its polyploid origin (Yang et al., 2022). Its maternal lineage is characterized by generalized / thick-lipped mouth polymorphism widely distributed among Torinae in the genera *Tor* and *Neolissochilus* (Hoang et al., 2015; Walton et al., 2017) – so the generalized/thick-lipped polymorphism is persistent within a Torinae lineage. Paternal lineage referred to contemporary genus *Cyprinion* is represented by scraping mouth phenotype (Coad, 2021). Hence, mouth polymorphism of the *Labeobarbus* is ancestrally heritable and re-evolved under particular ecological circumstances. Given this, the thick-lipped and scraping mouthed phenotypes of the *Labeobarbus* in the Sore River are seemingly ‘preadaptive’ upon emergence *de-novo*.

Modular-assembling genome of the *Labeobarbus* consisted of the different subgenomes (Yang et al., 2022) that is able to produce discrete mouth phenotypes inherent to parental lineages is formally still within ‘flexible stem hypothesis’. However, from another hand, it might be also in frame of ‘transporter process’ (Schluter & Conte, 2009; Marques et al., 2019; Martin & Richards, 2019) meaning that the adaptive alleles and genetic architectures differentiating each species within a rapid radiation are older than the radiation itself. The oldest paleorecords of the *Labeobarbus* in East Africa are dated by Late Miocene (Stewart & Murray, 2017). Some trophic specialists, for instance, with scraping phenotype are remarkably older than the recently emerged repeated radiations in the Ethiopian Highlands under consideration. Their age is dated as Pleistocene according to molecular clocks (Beshera et al., 2016). It suggests that similar mouth diversification might evolved repeatedly before the diversifications we observe in this study. Therefore, the genomes of the modern *Labeobarbus* might be rather experienced, i.e. may have the standing genetic variation contributing to the ecological speciation and diversification. Regardless which is the certain prerequisite for adaptive radiation of the *Labeobarbus* – ‘flexible stem’ or ‘transporter process’ (or their combination) – it might significantly facilitate the diversification.

### 4.3. Possible scenario of evolution of trophic radiation: from incipient to matured

Our results demonstrate that four riverine assemblages of the *Labeobarbus* in the Ethiopian Highlands being similar in the core mouth phenotypes and direction of trophic diversification are at various stages of the evolution. The number of mouth phenotypes together with subtypes and their ecologic functionality in the *Labeobarbus* varied significantly from river to river. We tried to range the cases from simplest to most matured based on the obtained results – diversity of ecomorphs/mouth phenotypes and their functionality (as estimated by the gut content and isotopic niche) within each radiation (Table 3; Figures 4-5).

**Table 3.**
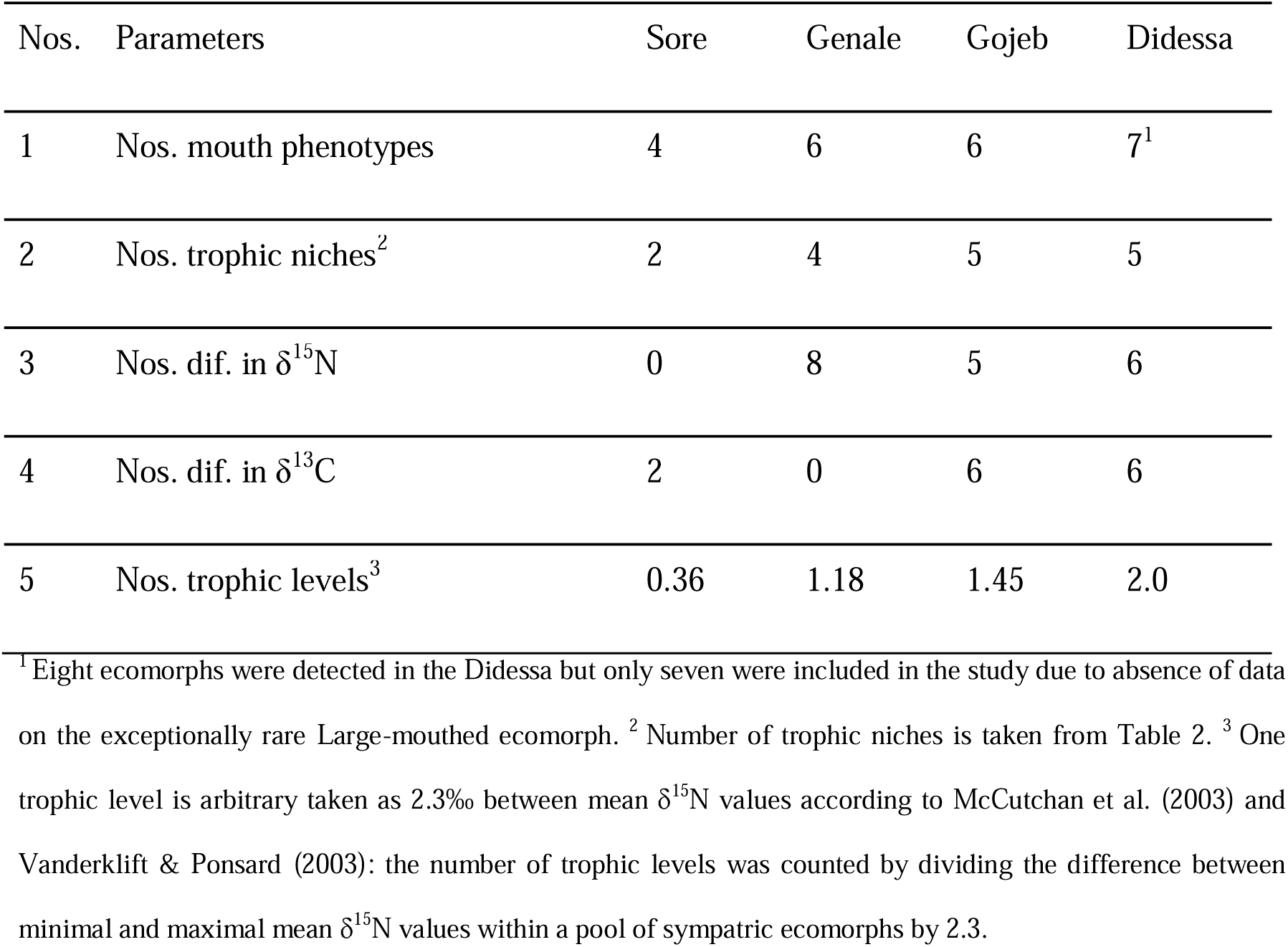
Ranking the diversifications by number of ecological niches and other aspects of trophic differentiation. Number of points in rows 3 and 4 corresponds to the number of significant differences in pairwise comparisons between sympatric ecomorphs (taken from Supplementary File S4).

The trophic diversifications could be ranged from the most undifferentiated in the Sore River via the Genale and Gojeb Rivers to the most diversified in the Didessa River (Table 3, Figure 7). Next stages of the trophic diversification can be provisionally outlined (see scheme on Figure 7):

1. Burst of mouth polymorphism. A set of core mouth phenotypes quickly emerges but their ecological functionalization is immature (preadaptive phenotypes). Morphological phenotypes are trophically irrelevant or there is a weak phenotype-trophic correlation (Liem’s paradox). At this stage phenotypic diversity strongly dominates over the ecological diversity - see the case of the Sore River (four phenotypes vs. two trophic niches).
2. Filling the trophic niches (ecological accommodation). Preadaptive phenotypes began to be functionalized by occupation of the trophic niches best matched to the mouth type. This process might be named as ecological accommodation of phenotypic trait – analogously to term ‘phenotypic accommodation’ sensu West-Eberhard (2003) The numbers of phenotypes and trophic niches are almost equal. Probably it is a very rapidly realizing process else it is hard to explain what are reasons to sustain non-functional mouth polymorphism.
3. Continued diversification within the core mouth phenotypes with corresponding increase of the ecological diversity. Upon filling the most evident trophic niches, a further sub-diversification occurs in some radiations. It might go by various scenarios - *via* ecological sub-specialization of certain phenotypes or *via* hybridization and functionalization of intermediate phenotypes. At matured stages of the diversification, the biotic interactions may serve as an additional ecological opportunity (‘diversity begets diversity’ – Martin & Richards, 2019). Significant increase of the populations size during morpho-ecological diversification (due to utilization of various trophic resources in a river) makes the piscivory strategy more reliable that may result in sub-diversification within this adaptive zone - like in the Didessa River (see also cascading speciation - Broderson et al., 2017; Bracewell et al., 2018).
4. A new diversification burst provoked by colonization of the novel environment might occur as exemplified from the lacustrine burst of the *Labeobarbus* in Lake Tana (Ethiopia) resulted in evolution of 15-16 species/ecomorphs (Nagelkerke et al., 1994; Mina et al., 1996; Sibbing, Nagelkerke, 1998; Nagelkerke et al., 2015; Beshera and Harris, 2023). It is still unknown whether Lake Tana radiation evolved from a single generalized ancestor or stemmed from already diversified set of riverine radiation.

**Figure 7.**
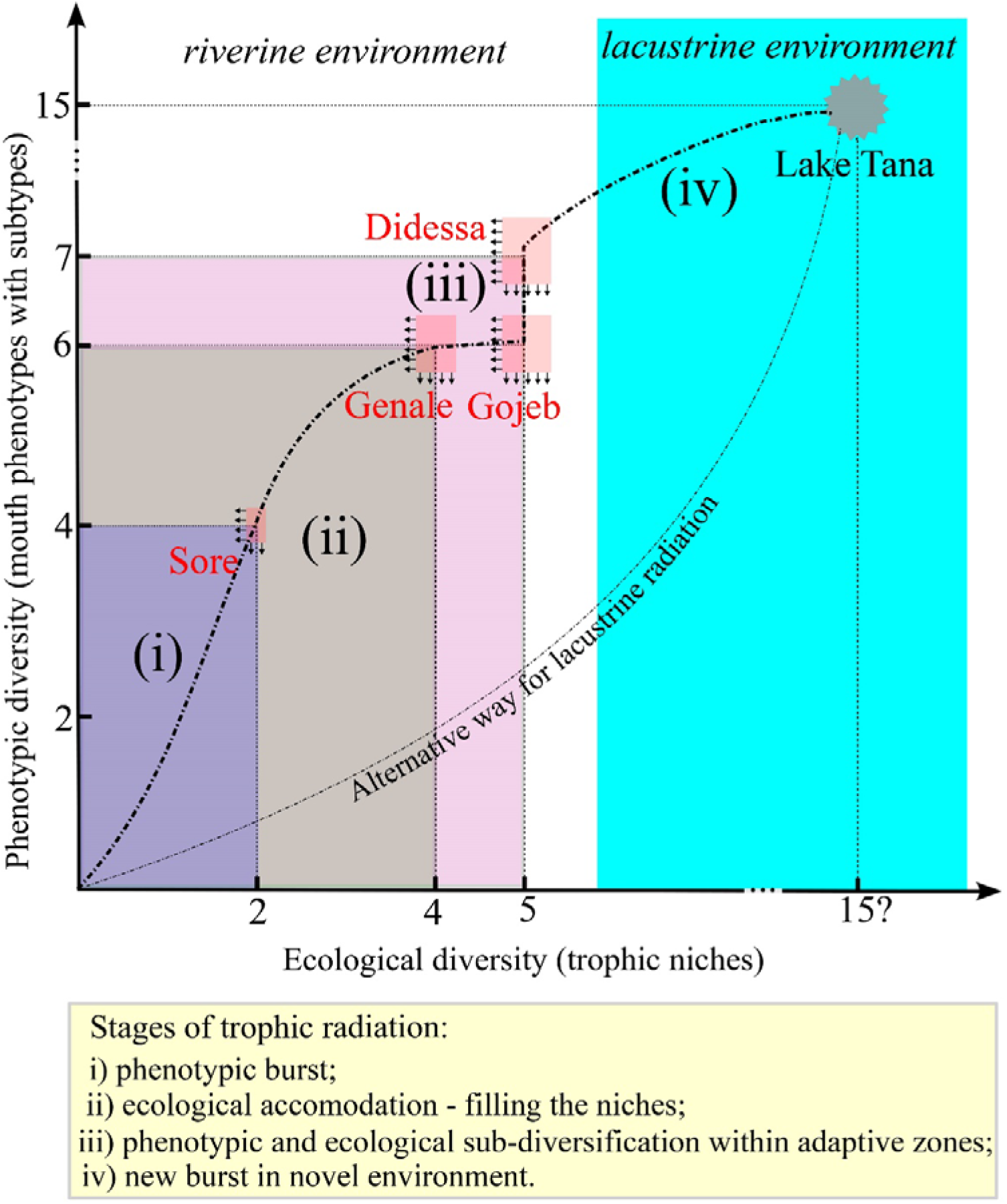
Schematic reconstruction of process of morpho-ecological diversification of the *Labeobarbus* in the Ethiopian Highlands. The number of little arrows at each box correspond to the numbers of mouth phenotypes (horizontal arrows) and numbers of trophic niches (vertical arrows).

The most immature diversification has been detected in the Sore River where a form (mouth phenotype) and its ecological function were largely decoupled. The ecologically non-functional or non-matured mouth polymorphism in the Sore River might be explained by several reasons. First, it might be due to a very young age of this radiation (at stage of incipient diversification) that is supported by low genetic diversity and absence of mtDNA haplotype sorting (Levin et al., 2020). Another reason refers to insufficient ecological opportunities in the river to realize the existing potential (mouth polymorphism). Heterogeneous and unpredictable environment in the rivers (flow regimes, turbidity, depth, water chemistry, temperature, food availability, etc.) may weaken the divergent selection necessary to drive specialization, instead favoring opportunistic ecological roles such as detritivory-omnivory (Jepsen and Winemiller, 2002; Burress et al., 2023). But the Sore River is a home for another bright adaptive radiation of hillstream cyprinid fish of the genus *Garra* that is represented by six genetically, morphologically and trophically divergent young species (Golubtsov et al., 2012; Levin et al., 2021a; Komarova et al., 2022). Possible overlapping in the trophic specializations (predator, periphyton feeding, benthic invertivore) with more advantageous cyprinid radiation in shared environment can acutely suppress or constrain the diversification process in the Sore population of the *Labeobarbus* resulting in freezing the non-functional mouth polymorphism stage for a while. Simultaneously, other studied riverine populations of the *Labeobarbus* have represented more advanced trophic diversification stages in absence of other sympatric radiations (as far as we know) and characterized by matured or ecologically functional mouth phenotypes.

## Conclusions

Using the polyploid African barbs of the genus *Labeobarbus* as model, we showed the parallel morpho-ecological diversifications being similar in this phenomenon with other prominent examples of adaptive radiation among vertebrate animals like cichlids (Seehausen, 2015; Torres-Dowdall & Meyer, 2021; Burress et al., 2023) and Caribbean *Anolis* (Losos, 2009; Stroud & Losos, 2020). The most striking difference from above-mentioned examples is the sharply preadaptive nature of inherited mouth polymorphism as a starter of radiation. Likely, the diversification of the *Labeobarbus* starts from the onset of genomic pre-existing templates of mouth polymorphism. The emerged mouth phenotypes are preadaptive but not (or weakly) functional at the incipient stage of trophic diversification demonstrating decoupled form and function. Enhancement of ecological functions of preadaptive phenotypes (ecological accommodation) increases upon filling the trophic niches and result in maturation of trophic specialization that is often continued by further sub-diversification of mouth types and sub-specialization within previously occupied adaptive zones. The preadaptive mouth polymorphism can be considered a key innovation of the *Labeobarbus* that makes the bridge between phenotypic diversification and ecological opportunities remarkably shorter that might accelerate diversification rates. It apparently explains why this lineage overcomes the obstacles in unstable riverine environments unfavored for adaptive radiations by other fish lineages.

## Acknowledgements

We are grateful to all members of the Joint Ethiopian–Russian Biological Expedition (JERBE), who participated in our field operations (S. E. Cherenkov, Y. Y. Dgebuadze, Genanaw Tesfaye, Fekadu Tefera, M. V. Mina, and I. S. Razgon), and especially to JERBE coordinator Dr. A. A. Darkov for his permanent and invaluable aid. We are grateful to S. E. Cherenkov for photographing the fish as well as to Y. Y. Dgebuadze and M. V. Mina for discussion of unpublished results. The study was supported by the Russian Science Foundation (grant no. 19-14-00218).

## Scientific field survey permission information

All fish samples were collected under the framework of Joint Ethiopian-Russian Biological Expedition using the permits of the Ministry of Innovation and Technology, Addis Ababa, Ethiopia provided to A.S.G and B.A.L. by National Fishery and Other Aquatic Life Research Center of the Ethiopian Institute of Agricultural Research - EIAR, Sebeta.

## Data availability

Raw isotopic data are available as Supplementary data.

## Supplementary data

Supplementary data to this article can be found online.

## Authors’ contributions

B.A.L. and A.S.G. conceived and designed the research, conducted field surveys and collected samples; A.S.K. and A.V.T. performed the stable isotope samples preparation and their analysis; A.S.K. performed diet analysis; All authors wrote the manuscript. All authors read and approved the final version of the manuscript.

